# Bone marrow adipose tissue is a unique adipose subtype with distinct roles in systemic glucose homeostasis

**DOI:** 10.1101/673129

**Authors:** Karla J. Suchacki, Adriana A.S. Tavares, Domenico Mattiucci, Erica L. Scheller, Giorgos Papanastasiou, Calum Gray, Matthew C. Sinton, Lynne E. Ramage, Wendy A. McDougald, Andrea Lovdel, Richard J. Sulston, Benjamin J. Thomas, Bonnie M. Nicholson, Amanda J. Drake, Carlos J. Alcaide-Corral, Diana Said, Antonella Poloni, Saverio Cinti, Gavin J. MacPherson, Marc R. Dweck, Jack P.M. Andrews, Michelle C. Williams, Robert J. Wallace, Edwin J.R. van Beek, Ormond A. MacDougald, Nicholas M. Morton, Roland H. Stimson, William P. Cawthorn

**Author notes:** ***Correspondence to***: William Cawthorn, University/BHF Centre for Cardiovascular Science, The Queen’s Medical Research Institute, Edinburgh BioQuarter, 47 Little France Crescent, Edinburgh, EH16 4TJ.

## Abstract

Bone marrow adipose tissue (BMAT) represents >10% of total adipose mass, yet unlike white or brown adipose tissues (WAT or BAT), its role in systemic metabolism remains unclear. Using transcriptomics, we reveal that BMAT is molecularly distinct to WAT but is not enriched for brown or beige adipocyte markers. Instead, pathway analysis indicated altered glucose metabolism and decreased insulin responsiveness in BMAT. We therefore tested these functions in mice and humans using positron emission tomography–computed tomography (PET/CT) with ^18^F-fluorodeoxyglucose, including establishing a new method for BMAT identification from clinical CT scans. This revealed that BMAT resists insulin- and cold-stimulated glucose uptake and is thus functionally distinct to WAT and BAT. However, BMAT displayed greater basal glucose uptake than axial bones or subcutaneous WAT, underscoring its potential to influence systemic glucose homeostasis. These PET/CT studies are the first to characterise BMAT function *in vivo* and identify BMAT as a distinct, major subtype of adipose tissue.

**HIGHLIGHTS:**

- Bone marrow adipose tissue (BMAT) is molecularly distinct to other adipose subtypes.
- BMAT is less insulin responsive than WAT and, unlike BAT, is not cold-responsive.
- Human BMAT has greater basal glucose uptake than axial bone or subcutaneous WAT.
- We establish a PET/CT method for BMAT localisation and functional analysis *in vivo*.

## INTRODUCTION

Adipose tissue plays a fundamental role in systemic energy homeostasis. In mammals it is typically classified into two major subtypes: white adipose tissue (WAT), which stores and releases energy and has diverse endocrine functions; and brown adipose tissue (BAT), which mediates adaptive thermogenesis (Cinti, 2018). Cold exposure and other stimuli also cause the emergence of brown-like adipocytes within WAT, typically referred to as “beige” adipocytes (Cinti, 2018). White, brown and beige adipocytes have attracted extensive research interest, owing largely to their roles and potential as therapeutic targets in metabolic diseases.

Adipocytes are also a major cell type within the bone marrow (BM), accounting for up to 70% of BM volume. Indeed, this BM adipose tissue (BMAT) can represent over 10% of total adipose tissue mass in healthy adults (Cawthorn et al., 2014). BMAT further accumulates in diverse physiological and clinical conditions, including aging, obesity, type 2 diabetes and osteoporosis, as well as therapeutic contexts such as radiotherapy or glucocorticoid treatment. Strikingly, BMAT also increases in states of caloric restriction (Scheller et al., 2016). These observations suggest that BMAT is distinct to WAT and BAT and might impact the pathogenesis of diverse diseases. However, unlike WAT and BAT, the role of BMAT in systemic energy homeostasis remains poorly understood.

The metabolic importance of WAT is highlighted in situations of both WAT excess (obesity) and deficiency (lipodystrophy), each of which leads to systemic metabolic dysfunction (Cinti, 2018). This largely reflects the key role of WAT as an insulin target tissue. Adipocyte-specific ablation of the insulin receptor in mice causes insulin resistance, glucose intolerance and dyslipidaemia (Qiang et al., 2016; Sakaguchi et al., 2017). Similar effects result from adipocytic deletion of *Slc2a4* (Glut4), the insulin-sensitive glucose transporter (Abel et al., 2001). Conversely, adipocyte-specific overexpression of *Slc2a4* reverses insulin resistance and diabetes in mice predisposed to diabetes (Carvalho et al., 2005). Thus, insulin-stimulated glucose uptake is fundamental to WAT’s role in systemic metabolic homeostasis.

In contrast to WAT, the defining function of BAT is in mediating adaptive thermogenesis via uncoupled respiration. This is driven by mitochondria expressing uncoupling protein-1 (UCP1), which are abundant in brown adipocytes. Cold exposure is the classical stimulator of BAT activity: cold-induced glucose uptake is a hallmark of BAT activation and can be quantified *in vivo* using positron emission tomography–computed tomography (PET/CT) with ^18^F-fluorodeoxyglucose (^18^F-FDG) (Cinti, 2018; Ramage et al., 2016a). Cold exposure or chronic sympathetic stimulation exert similar effects on beige adipocytes, and activation of brown or beige adipocytes can enhance energy expenditure (Cinti, 2018). Consequently, the past decade has seen extensive interest in activating BAT, or promoting beiging of WAT, to treat metabolic disease (Cinti, 2018).

Compared to WAT and BAT, study of BMAT has been relatively limited. However, given its abundance and clinical potential (Scheller et al., 2016), BMAT is now attracting increasing attention, with several studies beginning to investigate its metabolic properties. BM adipocytes (BMAds) have been proposed to exist in two broad subtypes: ‘constitutive’ BMAds (cBMAds) appear as contiguous groups of adipocytes that predominate at distal skeletal sites, whereas ‘regulated’ BMAds (rBMAds) occur interspersed with the haematopoietic BM in the proximal and axial skeleton (Craft et al., 2018). Both subtypes are morphologically similar to white adipocytes, with large unilocular lipid droplets; however, their lipid content differs, with cBMAds having a greater proportion of unsaturated fatty acids than rBMAds or white adipocytes (Scheller et al., 2016). Like white adipocytes, BMAds also produce adipokines such as leptin and adiponectin (Sulston and Cawthorn, 2016) and can release free fatty acids in response to lipolytic stimuli, albeit to a lesser extent than WAT (Scheller et al., 2018; Tran et al., 1981). This lipolysis resistance is more pronounced for rBMAds, underscoring the functional differences in BMAd subtypes.

Despite these advances in understanding of BMAT lipid metabolism, its insulin responsiveness and role in systemic glucose homeostasis is poorly understood. PET/CT studies have demonstrated glucose uptake into whole bones or BM of animal models and humans (Huovinen et al., 2016; Huovinen et al., 2014; Nishio et al., 2012; Zoch et al., 2016), but uptake specifically into BMAT has not previously been examined. Whether BMAT is BAT- or beige-like is also debated (Scheller et al., 2016). UCP-1 positive adipocytes have been noted in vertebral BM of a young mouse (Nishio et al., 2012) and as an incidental finding in one clinical case study (Chapman and Vega, 2017), but most studies find very low skeletal UCP-1 expression (Krings et al., 2012; Sulston et al., 2016). It has been suggested that BMAT is BAT-like, albeit based only on transcript expression from whole bones (Krings et al., 2012). Notably, no studies have fully investigated if BMAT has properties of BAT or beige fat *in vivo*. Together, it remains unclear if BMAT performs metabolic functions similar to WAT, BAT or beige adipose tissue.

Herein, we used transcriptomic analysis and ^18^F-FDG PET/CT to address these fundamental gaps in knowledge and thereby determine if, *in vivo*, BMAT has metabolic functions of WAT or BAT. Our studies in animal models and humans demonstrate that BMAT is transcriptionally and functionally distinct to WAT, BAT and beige adipose tissue, identifying BMAT as a unique class of adipose tissue. We show that BMAT has greater basal glucose uptake than WAT and establish methods for BMAT characterisation by PET/CT. Together, this knowledge underscores the potential for BMAT to influence metabolic homeostasis and sets a foundation for future research to reveal further roles of BMAT in normal physiology and disease.

## RESULTS

### BMAT is transcriptionally distinct to WAT, BAT and beige adipose tissues

The functional hallmarks of WAT, BAT and beige adipose tissue are reflected on a molecular level, with each class having distinct transcriptomic profiles and characteristic marker genes (Rosell et al., 2014; Svensson et al., 2011; Wu et al., 2012). Thus, to test if BMAT has distinct metabolic functions, we first compared the transcriptomes of whole BMAT and WAT from two cohorts of rabbits. Principle component analysis of both cohorts identified BMAT as a distinct depot compared to gonadal WAT (gWAT) and inguinal WAT (iWAT) (Fig. 1A); however, BMAT from either rabbit cohort was not uniformly enriched for markers of brown or beige adipocytes (Fig. 1B, S1A): although *SLC27A2* was significantly higher in BMAT than WAT from both cohorts, and *PPARGC1A* in BMAT from cohort 1, several other brown and/or beige markers were more highly expressed in WAT, while most such markers were not differentially expressed between BMAT and WAT in either cohort (Fig. 1B, S1A). Thus, the transcriptomic distinction with WAT is not a result of BMAT being more brown- or beige-like. Instead, gene set enrichment analysis (GSEA) highlighted the potential for BMAT to have altered glucose metabolism and decreased insulin responsiveness compared to WAT (Fig. 1C,1D, S1B).

**Figure 1 –.**
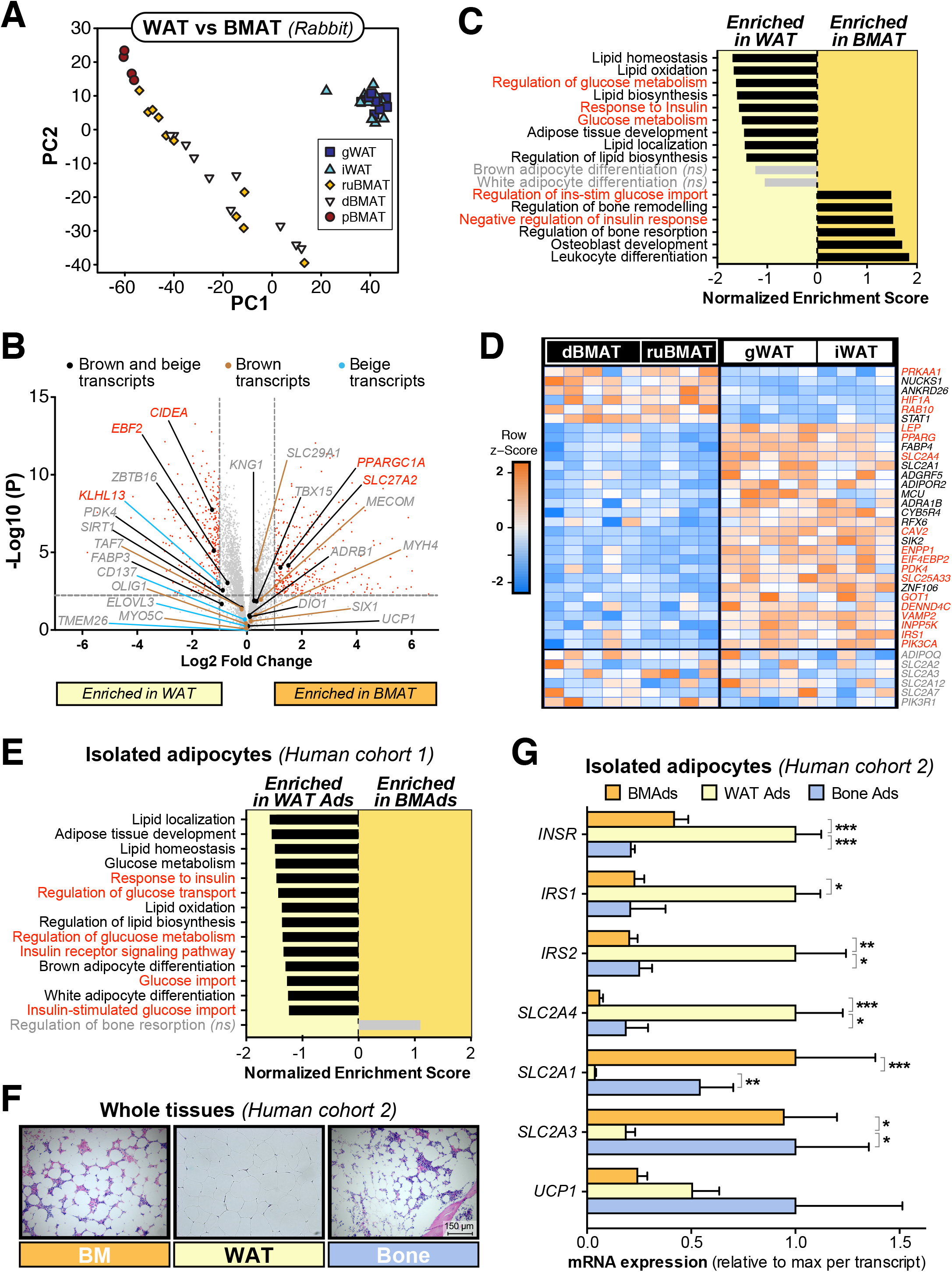
BMAT is transcriptionally distinct to white, brown and beige adipose tissues. (**A-D**) Transcriptional profiling of gonadal WAT, inguinal WAT, and whole BMAT isolated from the proximal tibia (pBMAT), distal tibia (dBMAT) or radius and ulna (ruBMAT) of two cohorts of rabbits. (A) Principal component analysis of both cohorts. (B-D) Volcano plots (B), GSEA (C) and heatmaps (D) of transcripts differentially expressed between BMAT (dBMAT + ruBMAT) and WAT (iWAT + gWAT) in rabbit cohort 1. In (B-E), red text indicates differentially expressed transcripts (B) or transcripts/pathways relating to glucose metabolism and/or insulin responsiveness (C-E); *ns* = not significant. (**E**) Transcriptional profiling of adipocytes isolated from femoral BM or subcutaneous WAT of humans. (**F**) Representative micrographs of H&E-stained sections of human femoral BM, subcutaneous WAT and trabecular bone; scale bar = 150 μm. (**G**) qPCR (G) of adipocytes isolated from tissues in (G). Data are mean ± SEM of the following numbers per group: BM Ads, n = 10; WAT Ads, n = 10; Bone Ads, n = 7 (*except IRS1, where n = 2 only*). For each transcript, significant differences between each cell type are indicated by * (*P* <0.05), ** (*P* <0.01) or *** (*P* <0.001). See also Figure S1 and Figure S2.

To determine if similar differences occur in humans, we next analysed the transcriptomes of adipocytes isolated from human femoral BMAT and subcutaneous WAT, which our previous analyses revealed to be globally distinct (Mattiucci et al., 2018). Consistent with our findings in rabbits, human BMAds were not enriched for brown or beige markers and had decreased expression of genes relating to glucose metabolism and insulin responsiveness (Fig. 1E, S2A, S2B). To further address this we next pursued targeted analysis of adipocytes isolated from BM and WAT of a second cohort of subjects; we also isolated adipocytes from trabecular bone (Bone Ads) to assess potential site-specific differences in BMAd function (Craft et al., 2018). Adipocyte purity was confirmed histologically (data not shown) and by qPCR (Fig. S2C). *In situ*, these BM and bone adipocytes resembled unilocular white adipocytes (Fig. 1F); however, qPCR revealed significant differences in transcript expression of *INSR, IRS1, IRS2, SLC2A4, SLC2A1* and *SLC2A3* between WAT adipocytes and those from BM or bone (Fig. 1G). Notably, compared to white adipocytes, each BMAd subtype had decreased *SLC2A4* and increased *SLC2A1* and *SLC2A3*, suggesting that BMAds may have higher basal glucose uptake that is less insulin responsive. In contrast, there were no differences in expression of *UCP1*, and most other brown or beige adipocyte markers were not enriched in either BMAd subtype (Fig. 1G, S2D).

Taken together, these data demonstrate that BMAds in animal models and humans are transcriptionally distinct to white, brown and beige adipocytes, and suggest altered roles in systemic glucose homeostasis and insulin responsiveness.

### Insulin treatment in mice does not induce glucose uptake in BMAT

To test the metabolic functions of BMAT *in vivo* we used ^18^F-FDG PET/CT in mice to determine if, like WAT, BMAT is insulin-responsive. As expected, insulin decreased blood glucose (Fig. 2A) and increased ^18^F-FDG uptake in the heart, iWAT and gWAT (Fig. 2B, C and F). To assess ^18^F-FDG uptake separately within bone and BMAT, we first applied thresholding to the PET/CT data to separate bone from BM based on their different tissue densities (*data not shown*). This revealed that insulin significantly increased ^18^F-FDG uptake in femoral bone, whereas humoral bone uptake decreased; uptake in proximal or distal tibial bone was unaffected (Fig. 2F). To assess BMAT-specific ^18^F-FDG uptake we took advantage of the regional differences in BMAT content around the mouse skeleton. Thus, adipocytes comprise only a small percentage of total BM volume of humeri, femurs and proximal tibiae, but predominate in distal tibiae (Fig. 2E). To address the contribution of BMAT to skeletal ^18^F-FDG uptake, we therefore quantified ^18^F-FDG in a bone-region-specific manner to distinguish between areas of low BMAT (humerus, femur, proximal tibia) and high BMAT (distal tibia). This revealed that insulin did not significantly affect glucose uptake in any of the BM regions analysed (Fig. 2F). Thus, compared to WAT, BM and BMAT resist insulin-stimulated glucose uptake.

**Figure 2 –.**
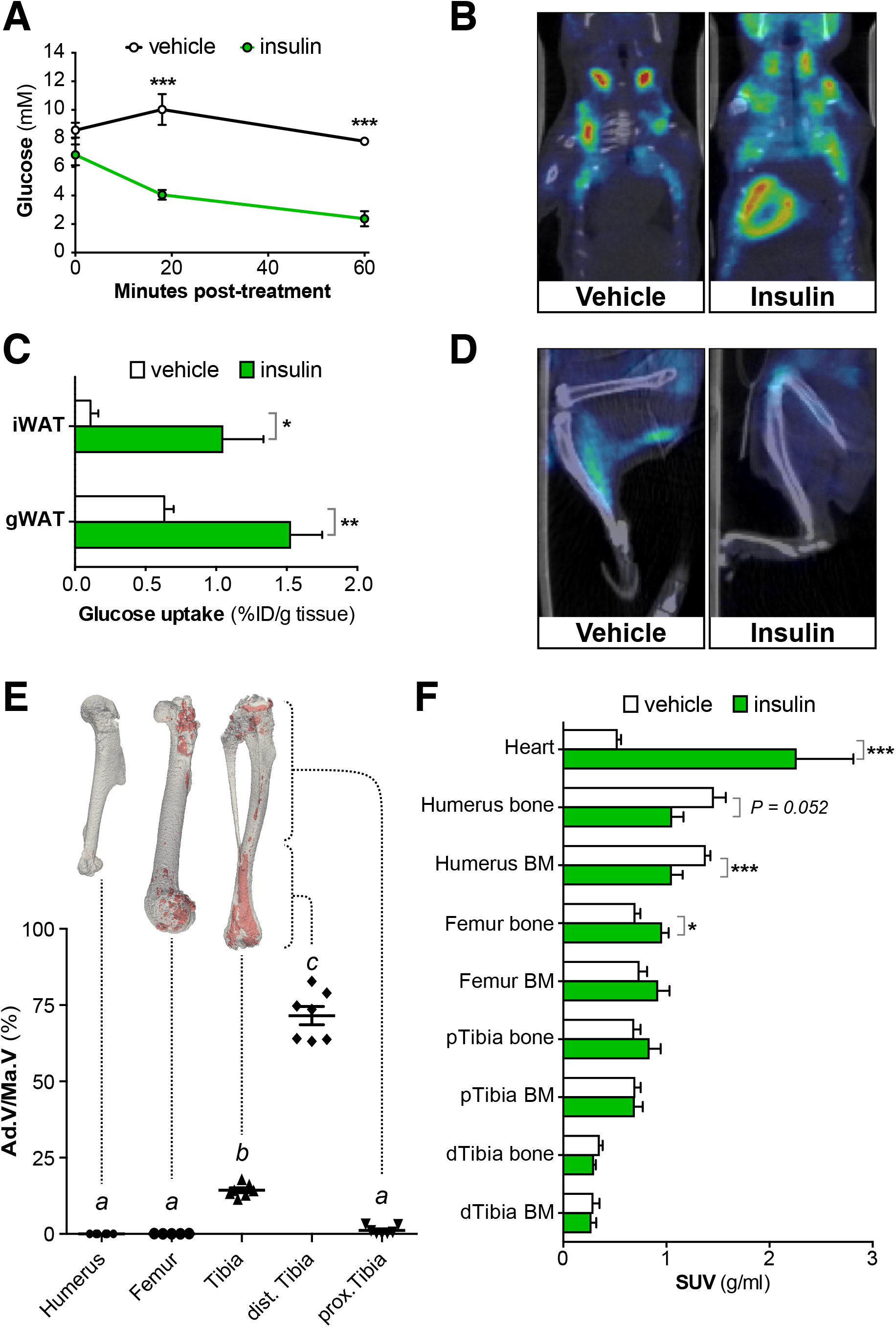
Insulin treatment in mice does not induce glucose uptake in BMAT. Insulin-stimulated glucose uptake was assessed by PET/CT. (**A**) Blood glucose post-insulin or vehicle. (**B,D**) Representative PET/CT images of the torso (B) or legs (D) of vehicle- and insulin-treated mice; some ^18^F-FDG uptake into skeletal muscle is evident in the image of the vehicle-treated mouse (D), possibly resulting from physical activity. (**C**) Gamma counts of ^18^F-FDG uptake in iWAT and gWAT, shown as *%* injected dose per g tissue (%ID/g). (**E**) BMAT analysis by osmium tetroxide staining. BMAT is shown in red in representative μCT reconstructions and quantified as adipose volume relative to total BM volume (Ad.V/Ma.V). (**F**) ^18^F-FDG uptake in the indicated tissues was determined from PET/CT scans. Data are presented as mean ± SEM of 5-6 mice (A,C,F) or 5-7 mice (E). Significant differences between control and insulin-treated samples are indicated by * (*P* <0.05), ** (*P* <0.01) or *** (*P* <0.001). In (E), groups do not significantly differ if they share the same letter.

### Cold exposure in mice does not induce glucose uptake or beiging in BMAT

To test if BMAT is BAT- or beige-like *in vivo*, we next analysed ^18^F-FDG uptake following acute or chronic cold exposure in mice (Fig. S3A). Acute (4 h) or chronic cold (72 h) increased energy expenditure without causing weight loss or hypoglycaemia (Fig. S3B-D), likely due to increased food consumption in chronic cold mice (Fig. S3E). BAT ^18^F-FDG uptake increased after either duration of cold (Fig. 3A-C). Chronic cold also increased iWAT ^18^F-FDG uptake, suggesting beiging of this depot (Fig. 3C). However, neither acute nor chronic cold exposure increased ^18^F-FDG uptake into bone or BM (Fig. 3B). Indeed, cold exposure decreased ^18^F-FDG uptake into distal tibial BMAT, highlighting fundamental differences with iWAT and BAT. Cold exposure also decreased BAT lipid content and promoted beiging of iWAT, as indicated by formation of multilocular adipocytes, but these effects did not occur in BMAT (Fig. 3D). Consistent with this, cold exposure induced brown and beige transcripts in BAT and iWAT, but not within bone (Fig. 3E-G, S3F-H). Housing control mice at room temperature (22 °C) might have caused a mild cold stress, preventing detection of further beiging at 4 °C; however, even when compared to mice at thermoneutrality (28 °C), cold exposure did not induce a beiging response within bones (Fig. S3I). These results show that, *in vivo*, BMAT is functionally distinct to brown and beige adipose tissues.

**Figure 3 –.**
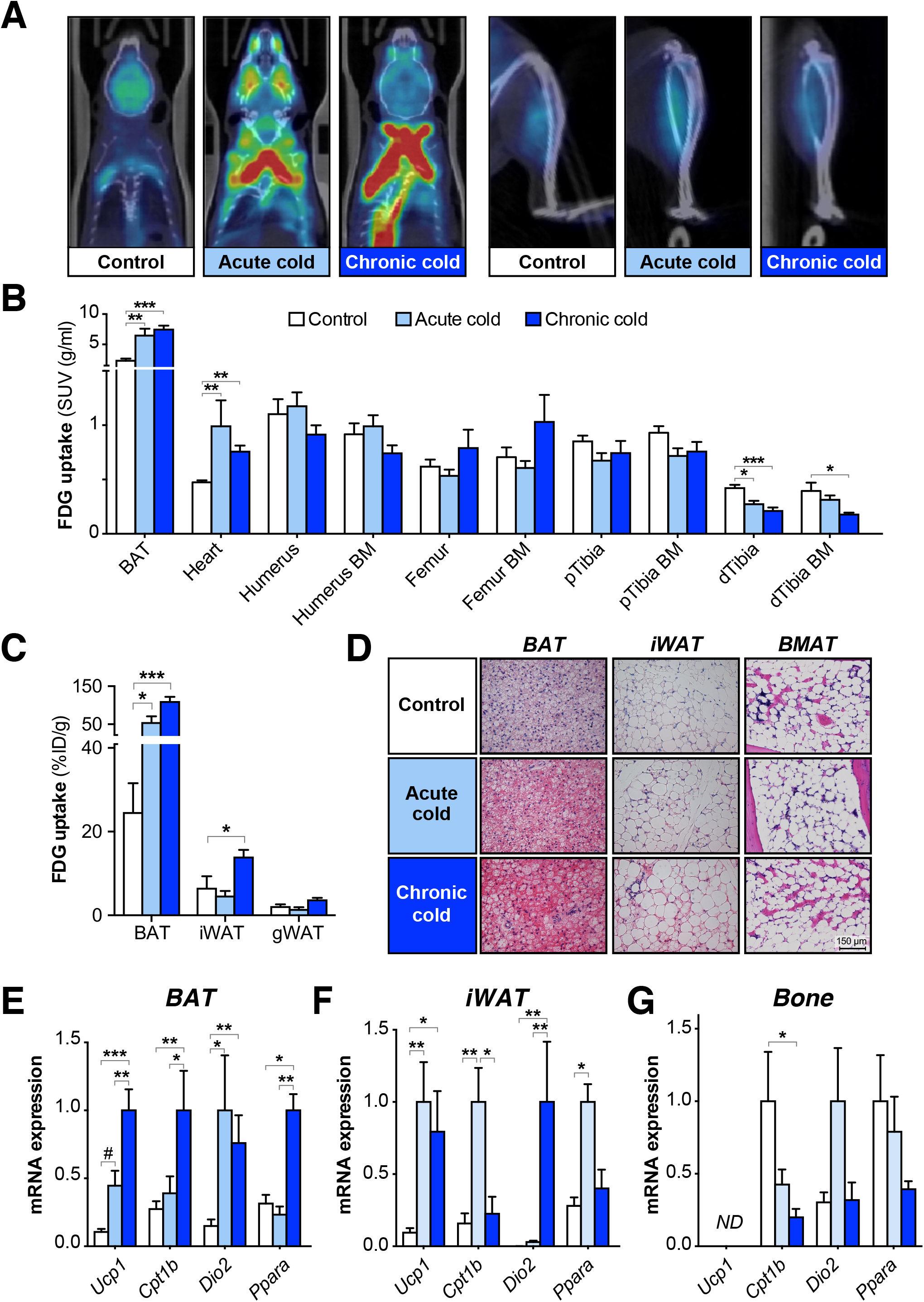
Cold exposure does not induce glucose uptake or beiging in BMAT. Cold-induced glucose uptake was assessed by PET/CT, as described in Figure S3A. (**A**) Representative PET/CT images of control, acute and chronic cold mice show increased ^18^F-FDG uptake in BAT but not tibiae; some ^18^F-FDG is evident in skeletal muscle of each group. (**B,C**) ^18^F-FDG uptake in the indicated tissues was determined by PMOD analysis of PET/CT scans (B) or gamma counting (C). (**D**) Representative micrographs of H&E-stained tissues, showing that cold exposure decreases lipid content in BAT and promotes beiging of iWAT, but these effects do not occur in BMAT; scale bar = 150 μm. (**E-G**) Cold exposure induces brown and beige adipocyte transcripts in BAT and iWAT, but not in whole bones. *ND =* not detectable. Data in (B-C) and (E-G) are shown as mean ± SEM of 7-8 mice per group. Significant differences between groups are indicated by # (*P* <0.01), * (*P* <0.05), ** (*P* <0.01) or *** (*P* <0.001). See also Figure S3.

### CT-based identification of BMAT in humans

We next tested if these distinct metabolic properties extend to BMAT in humans. First, we established a method to identify BMAT from clinical PET/CT scans. To determine Hounsfield Units (HU) for BMAT, we identified BMAT-rich and BMAT-deficient BM regions by magnetic resonance imaging (MRI). This revealed that sternal BM is BMAT-enriched while vertebral BM is BMAT-deficient (Fig. 4A); WAT was also analysed as an adipose-rich control region. We then co-registered the MRI data with paired CT scans of the same subjects (Fig. 4A). This revealed a distinct HU distribution for BMAT-rich sternal BM, intermediate between WAT and red marrow (RM) of BMAT-deficient vertebrae (Fig. 4B).

**Figure 4 –.**
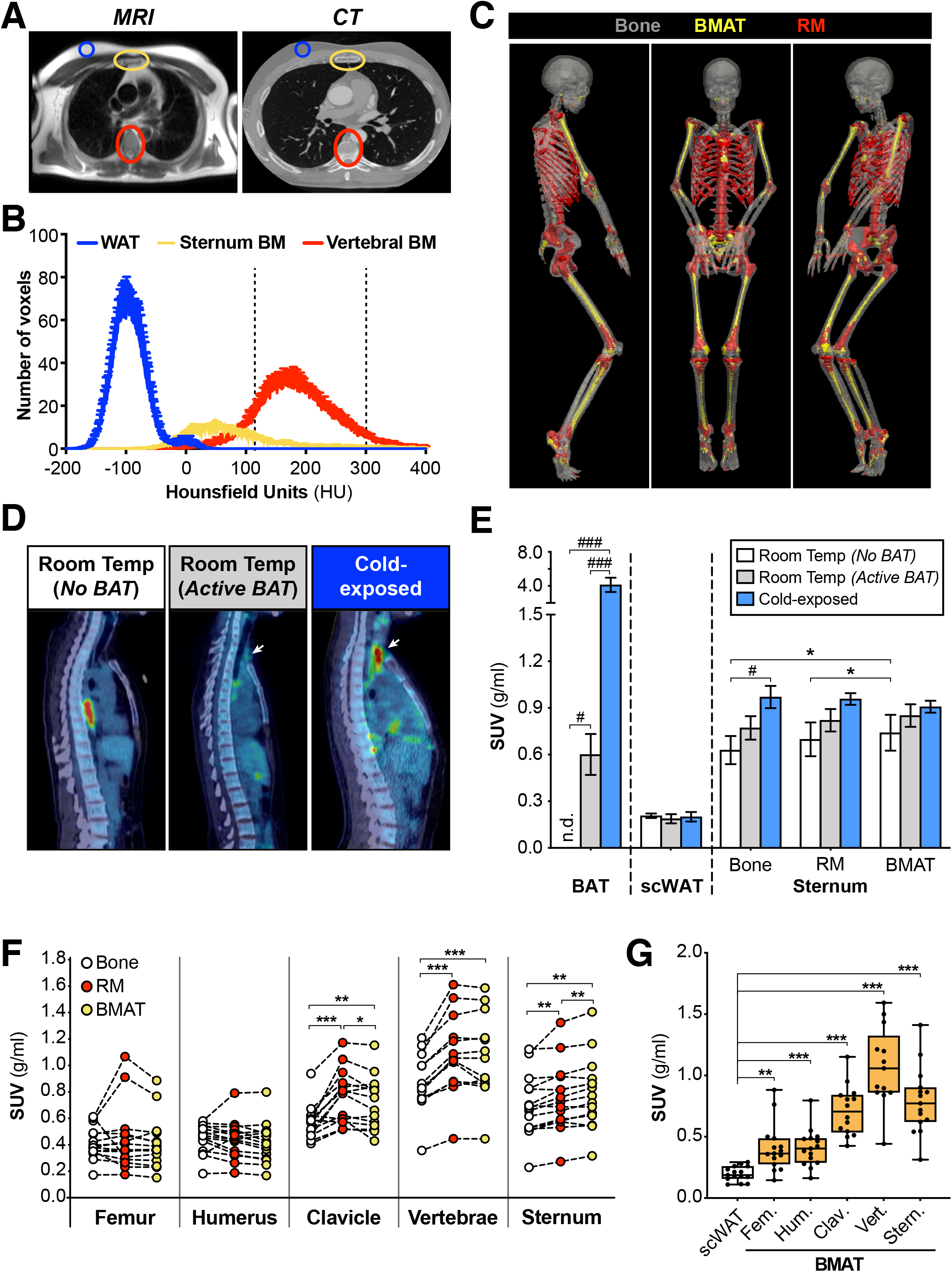
Human BMAT is functionally distinct to BAT and is a major site of basal glucose uptake. (**A**) Representative MRI (HASTE) and CT images from one subject. (**B**) HU distribution of scWAT, BMAT-rich BM (sternum) and BMAT-deficient BM (vertebrae). Data are mean ± SEM (n = 33). Thresholds diagnostic for BMAT (<115) and RM (115-300) are indicated by dashed lines. (**C**) CT images of a 32-year-old subject, highlighting BMAT or RM identified using the diagnostic thresholds in (B). Tibiae are shown for completeness but were not present in any other available CT scans. (**D,E**) PET/CT analysis of ^18^F-FDG uptake in *No BAT, Active BAT* and cold-exposed (*Cold*) subjects. Representative PET/CT scans in (D) highlight the BM cavities of the vertebrae and sternum; arrows indicate ^18^F-FDG uptake in supraclavicular BAT. (**F,G**) ^18^F-FDG uptake in bone tissue, RM and BMAT (F), or BMAT and scWAT (G), of room-temperature subjects (*No Bat* and *Active BAT* groups); Fem. = femur, Hum. = humerus, Clav. = clavicle, Vert. = vertebrae, Stern. = sternum. Data are shown as mean ± SEM (E), paired individual values (F) or box-and-whisker plots (G) of 8 (*No BAT*) or 7 (7 *Active BAT, Cold*) subjects per group. Significant differences between bone, RM and BMAT are indicated by * (*P* <0.05), ** (*P* <0.01) or *** (*P* <0.001). Significant differences between *No BAT, Active BAT* and *Cold* groups are indicated by # (*P* <0.05) or ### (*P* <0.001). See also Figure S4.

Using these distinct HU distributions, we generated a receiver operating characteristic (ROC) curve to identify optimal diagnostic HU thresholds to distinguish BMAT from RM (Fig. S4A). This revealed that BMAT-rich BM has HU <115, whereas RM is mostly within 115-300 HU (Fig. 4B); bone was defined as >300 HU. To test the validity of these thresholds we applied them to clinical CT data to determine BMAT volume as *%* BM volume. We found that BMAT predominated in the arms, legs and sternum but was markedly lower in the clavicle, ribs and vertebrae (Fig. 4C, Fig. S4B). Moreover, %BMAT showed age-associated increases in the axial skeleton but not in long bones (Fig. S4B). These data are consistent with previous studies showing that BMAT predominates in the long bones by early adulthood but continues to accumulate in axial bones beyond 60 years of age (Baum et al., 2018; Kricun, 1985; Schraml et al., 2015). Together, this supports the validity of our CT thresholds for BMAT identification in humans.

### Human BMAT is functionally distinct to BAT and is a major site of basal glucose uptake

We then applied these thresholds to human co-registered PET/CT data to assess ^18^F-FDG uptake in BMAT, RM and bone. To test if BMAT is BAT-like we first compared BMAT ^18^F-FDG uptake between three groups: subjects with no detectable supraclavicular BAT at room temperature (*No BAT*), subjects with active BAT at room temperature (*Active BAT*), and cold-exposed subjects (16 °C for 2 h; *Cold*). PET/CT confirmed BAT ^18^F-FDG uptake in the latter two groups but not in the *No BAT* group (Fig. 4D-E, Fig. S4C). Cold exposure did not alter ^18^F-FDG uptake in scWAT but was associated with increased uptake in sternal and clavicular bone tissue; however, these were the only skeletal sites at which ^18^F-FDG uptake significantly differed between the *No BAT, Active BAT* and *Cold* subjects (Fig. 4E, Fig. S4D). Indeed, the *Active BAT* and *Cold* subjects did not have increased glucose uptake in BMAT or RM of any bones analysed (Fig. 4E, Fig. S4D). Thus, consistent with our findings in mice, BMAT glucose uptake in humans is not cold-responsive.

Our previous human PET/CT studies revealed that glucocorticoids acutely activate BAT (Ramage et al., 2016a). Glucocorticoids also promote BMAT accumulation, demonstrating that BMAT can be glucocorticoid-responsive (Scheller et al., 2016). Thus, to further test if BMAT shares properties of BAT, we analysed PET/CT data from previously reported subjects (Ramage et al., 2016a) to determine if glucocorticoids also influence glucose uptake in human BMAT. We found that prednisolone significantly influenced ^18^F-FDG uptake only in vertebrae, in which there was a trend for increased uptake into RM but not BMAT or bone (Fig. S4E). However, prednisolone did not influence ^18^F-FDG uptake at any other site. Thus, unlike BAT, BMAT glucose uptake is not glucocorticoid-responsive.

The above findings confirm that, in humans, BMAT is functionally distinct to BAT. However, while analyzing these data, two other phenomena became apparent. Firstly, within axial bones of each subject, BMAT had significantly higher glucose uptake than bone (Fig. 4F). In the sternum, glucose uptake was also greater in BMAT than in RM (Fig. 4F). Secondly, BMAT at each skeletal site had higher glucose uptake than scWAT (Fig. 4G). Thus, despite being unresponsive to insulin or activators of BAT, BMAT has high basal glucose uptake, highlighting its potential to influence systemic glucose homeostasis.

## DISCUSSION

Unlike WAT and BAT, the role of BMAT in systemic energy metabolism is poorly understood. Previous studies have shed some light on BMAT lipid metabolism *in vivo* (Scheller et al., 2018; Tran et al., 1981), and PET/CT has been used to assess glucose uptake into bones or BM (Huovinen et al., 2016; Huovinen et al., 2014; Nishio et al., 2012; Zoch et al., 2016); however, our study is the first to characterise *in vivo* glucose metabolism specifically in BMAT. Our data provide key insights into how BMAT compares to WAT and BAT; reveal new site-specific differences in BMAT characteristics; and identify BMAT as a major site of skeletal glucose disposal. Moreover, we establish a method for BMAT identification and analysis by PET/CT that will open new avenues for future study of BMAT function.

We show, for the first time, that, compared to WAT, BMAT resists insulin-stimulated glucose uptake. This is supported not only by PET/CT of mouse distal tibial BMAT, but also by the transcriptional profiles of rabbit and human BMAT from other skeletal sites. This conclusion seemingly contrasts with findings elsewhere. For example, adipocyte-specific ablation of *Insr* in mice decreases BMAd size (Qiang et al., 2016), suggesting a role for insulin in BMAd lipogenesis; however, it is unclear if this is through *de novo* lipogenesis from glucose, or via insulin regulating uptake and esterification of fatty acids. In humans, Huovinen *et al* used PET/CT to assess BM ^18^F-FDG uptake during hyperinsulinaemic euglycemic clamp, concluding that whole BM may be insulin-responsive (Huovinen et al., 2016). Thus, one possibility is that insulin can stimulate BMAT glucose uptake under hyperinsulinaemic conditions. However, unlike our work, Huovinen *et al* did not distinguish BMAT-rich from BMAT-deficient BM, nor did they use a vehicle control to confirm if BM ^18^F-FDG uptake is genuinely insulin-responsive. Indeed, microarrays show that *SLC2A4* and *IRS1* expression is negligible in human BM (Dezso et al., 2008), while *Slc2a4* and *Irs1* are markedly lower in BM than in WAT or muscle of mice (Thorrez et al., 2008). More recent microarrays show that *Slc2a4, Insr, Irs1* and *Irs2* are lower in BMAds than epididymal white adipocytes of mice (Liu et al., 2011). These data are strikingly consistent with our results for transcript expression in rabbits and humans (Fig. 1, Fig. S1–2) and further support the conclusion that, compared to WAT, BMAT resists insulin-stimulated glucose uptake.

In addition to BMAT, we also found that insulin responsiveness varies among different bones: in insulin-treated mice, bone glucose uptake increases in femurs, decreases in humeri and is unaltered in tibiae. In contrast, Zoch *et al* report that insulin stimulates ^18^F-FDG uptake into whole femurs and tibiae (Zoch et al., 2016). This discrepancy may relate to technical differences: Zoch *et al* analysed whole bones (including BM) of anesthetised mice, whereas we distinguished between bone and BM and avoided anaesthesia. It is unclear why insulin is associated with decreased glucose uptake in humeral bone and BM; this is unlikely to be a technical issue given that we see expected insulin-stimulated glucose uptake in the heart, WAT and femur. Thus, the lack of increases in humeri and tibiae suggests that there are site-specific differences in skeletal insulin responsiveness.

Another major finding is that BMAT is molecularly and functionally distinct to brown and beige adipose tissues, both for cBMAT of mice and rabbits, and for rBMAT of humans at multiple skeletal sites. These molecular distinctions are consistent with several other studies. We and others previously found that tibial *Ucp1* expression is over 10,000-fold lower than in BAT (Krings et al., 2012; Sulston et al., 2016), consistent with our present finding that *Ucp1* is undetectable in whole mouse bones. Similarly, microarrays show that *UCP1* is not enriched in whole BM of mice or humans (Dezso et al., 2008; Thorrez et al., 2008), nor is it greater in BMAds vs white adipocytes of mice (Liu et al., 2011). Moreover, BMAd progenitors are more white-like than brown-like and do not express brown adipocyte markers after adipogenesis *in vitro* (Ambrosi et al., 2017). However, despite these diverse lines of evidence to the contrary, the concept that BMAT may be BAT- or beige-like has persisted. Thus, our *in vivo* functional analyses of mice and humans are a key advance because they confirm that cold exposure does not induce glucose uptake or beiging in BMAT. This demonstrates, conclusively, that BMAT is not BAT- or beige-like.

Our glucocorticoid studies provide further insights. Unlike in BAT, acute glucocorticoid treatment in humans does not stimulate glucose uptake in BMAT; however, it does influence uptake across lumbar vertebrae, with a trend for increases in RM (Fig. S4E). It is notable that this occurs only in vertebrae, because these are also the bones in which glucocorticoids drive the greatest increases in fracture risk (Briot and Roux, 2015). This raises the possibility that glucocorticoids modulate BM and bone metabolism in a site-specific manner and that these metabolic effects contribute to glucocorticoid-induced osteoporosis. Future studies using different doses and durations of glucocorticoids would further elucidate their ability to modulate metabolism of RM, BMAT and bone, and whether this influences glucocorticoid-induced osteoporosis.

Although BMAT glucose uptake is not stimulated by insulin at physiological concentrations, cold exposure or glucocorticoids, a major finding is that BMAT in humans has high basal glucose uptake, exceeding that of WAT and greater than that for bone or RM in the axial skeleton. Superficially, this seems at odds with two studies reporting that BM ^18^F-FDG uptake correlates inversely with BM fat content (Huovinen et al., 2014; Schraml et al., 2015); however, on further consideration, it is clear that these findings are not inconsistent with ours. Indeed, we show that axial bones have less BMAT but greater BM ^18^F-FDG uptake than humeri or femurs, mirroring these and other previous reports of ^18^F-FDG uptake in whole BM (Huovinen et al., 2016). Importantly, unlike our approach, no previous studies have distinguished ^18^F-FDG uptake between RM and BMAT specifically. Thus, a unique advance of our work is the finding that, in axial bones, BMAT glucose uptake is greater than in bone and similar or greater than in RM. This is particularly notable given that both BM and bone are sites of high glucose uptake, capable of exceeding levels observed in WAT or skeletal muscle (Huovinen et al., 2016; Huovinen et al., 2014; Zoch et al., 2016) (Fig. 4, Fig. S4). Indeed, bone glucose uptake is required for normal metabolic function (Li et al., 2016). Together, these observations support the conclusion that BMAT may influence systemic glucose homeostasis.

We also reveal that BMAT glucose uptake varies at different skeletal sites, generally being greater in axial BMAT compared to BMAT in long bones. This is consistent with depot-dependent differences in other BMAT characteristics and, broadly, with the concept that BMAT exists in regulated and constitutive subtypes (Craft et al., 2018). However, while axial BMAT has higher glucose uptake, BMAT volume in peripheral bones is typically far higher (Fig. 4, Fig. S4) (Kricun, 1985). Thus, the systemic metabolic influence of axial BMAT may be greater in conditions such as ageing, obesity, osteoporosis or caloric restriction, in which axial BMAT accumulates (Scheller et al., 2016).

Why does BMAT have such high basal glucose uptake? This may result from BMAds having high expression of *SLC2A1* and/or *SLC2A3* (Fig. 1), a finding supported by previous microarray studies (Liu et al., 2011). Indeed, among numerous human tissues, *SLC2A3* expression is highest in BM (Dezso et al., 2008), while *Slc2a3* is also greater in BM than in WAT of mice (Thorrez et al., 2008). Mouse BMAds also have a dense mitochondrial network (Robles et al., 2018) and, by electron microscopy, we found that mitochondria are also abundant in human BMAds (S. Cinti, personal communication). This supports the conclusion that BMAds are metabolically active, which may further explain their high basal glucose uptake.

Finally, we have developed a method to identify BMAT from CT scans, allowing its functional analysis by PET/CT. At least one other study has used HU thresholding to try to distinguish BMAT-enriched vs BMAT-deficient BM (Rantalainen et al., 2013), but our method is more comprehensive because we directly compared paired MRI and CT scans to identify the optimal BMAT HU thresholds. The finding that the sternum is BMAT-rich was unexpected as this contrasts with most other axial bones; however, it is consistent with adipogenic progenitors being readily detectible within sternal BM (Ambrosi et al., 2017). Otherwise, our method identifies site- and age-dependent differences in RM and BMAT that are in full agreement with previous studies (Baum et al., 2018; Kricun, 1985; Schraml et al., 2015). Applying our PET/CT approach to other clinical and preclinical studies, including retrospectively, therefore holds great promise to reveal further physiological and pathological roles of BMAT. Importantly, the diversity of PET tracers could extend such studies far beyond glucose metabolism, allowing many other functions of BMAT to be addressed.

In summary, this study is the first to dissect BMAT glucose metabolism *in vivo* and identifies BMAT as a distinct, major subtype of adipose tissue.

## ACKNOWLEDGEMENTS

This work was supported by grants from the Medical Research Council (MR/M021394/1 to W.P.C.; MR/K010271/1 to R.H.S.), the National Institutes of Health (R01 DK62876 and R24 DK092759 to O.A.M.; K99-DE024178 to E.L.S.; and P30 DK089503 to the Michigan Nutrition Obesity Research Center), the Wellcome Trust-University of Edinburgh Institutional Strategic Support Fund (to W.P.C. and K.J.S.), and the British Heart Foundation (4-year BHF PhD Studentship to R.J.S., B.J.T., M.C.S. and B.M.N; BHF CoRE Bioinformatics Grant to W.P.C.; BHF CoRE grant to A.J.D.). W.P.C. is further supported by a Chancellor’s Fellowship from the University of Edinburgh. A.A.S.T was funded by the British Heart Foundation (RG/16/10/32375). E.J.R.v.B is supported by SINAPSE (the Scottish Imaging Network). We are grateful to the British Heart Foundation for providing funding towards establishment of the Edinburgh Preclinical PET/CT laboratory (RE/13/3/30183), and to NHS Research Scotland (NRS) for financial support of the Edinburgh Clinical Research Facility. R.H.S and M.C.W. are supported by The Chief Scientist Office of the Scottish Government (SCAT/17/02 to R.H.S.; PCL/17/04 to M.C.W.). J.P.M.A is supported by BHF Clinical Research Training Fellowship no. FS/17/51/33096. We are grateful to John Henderson (BVS, University of Edinburgh) for support with mouse husbandry; Anish K. Amin (Department of Orthopaedic Surgery, Royal Infirmary Edinburgh), Beena Polouse and Frank Morrow (Edinburgh Clinical Research Facility,) for help with human studies; Tashfeen Walton and Christophe Lucatelli (Edinburgh Imaging, University of Edinburgh) for radiotracer production; Robert K. Semple (Centre for Cardiovascular Science, University of Edinburgh) for critical feedback on this manuscript; and to staff at the University of Michigan microarray core facility for processing of rabbit microarray data.

## AUTHOR CONTRIBUTIONS (based on CRediT taxononmy)

***Conceptualisation***, K.J.S. and W.P.C.; ***Methodology***, K.J.S., A.A.S.T, E.L.S., G.P., C.G., L.E.R., W.A.M., A.J.D., A.P., S.C., R.J.W., O.A.M., N.M.M, R.H.S. and W.P.C.; ***Investigation***, K.J.S., D.M., E.L.S., L.E.R., G.P., C.G., A.L., R.J.S., B.J.T., M.C.S., B.M.N., C.J.A., D.S., G.J.M., M.R.D., J.P.M.A., M.C.W., R.J.W., E.J.R.v.B., R.H.S. and W.P.C.; ***Formal Analysis***, K.J.S., A.A.S.T, D.M., E.L.S., G.P., C.G., W.A.M., J.P.M.A., M.C.W., R.J.W., E.J.R.v.B., N.M.M., R.H.S. and W.P.C.; ***Resources***, A.A.S.T, A.P., S.C., G.J.M., M.R.D., J.P.M.A., M.C.W., E.J.R.v.B., N.M.M., R.H.S.; Writing – Original Draft, K.J.S. and W.P.C.; ***Writing – Review & Editing***, K.J.S., A.A.S.T, E.L.S., G.P., W.A.M., A.J.D., S.C., M.C.W., R.J.W., E.J.R.v.B., O.A.M., R.H.S, W.P.C.; ***Visualisation***, K.J.S. and W.P.C.; Supervision, A.J.D., A.P., M.R.D., O.A.M., N.M.M., R.H.S. and W.P.C.; ***Funding Acquisition***, A.J.D., O.A.M., N.M.M., R.H.S. and W.P.C.

## METHODS

### Table of key resources

| REAGENT or RESOURCE | SOURCE | IDENTIFIER |
| --- | --- | --- |
| BIOLOGICAL SAMPLES |  |  |
| Human bone marrow (*cohort 1*) | Orthopedic and Traumatology Department, Ospedali Riuniti, Ancona, Italy | N/A |
| Human white adipose tissue (*cohort 1*) | Hepatobiliary and Abdominal Transplantation Surgery, Department of Experimental and Clinical Medicine, Università Politecnica delle Marche, Ancona, Italy | N/A |
| Human bone marrow and white adipose tissue (*cohort 2*) | Edinburgh Adipose Tissue Biobank | N/A |
| CHEMICALS |  |  |
| Collagenase Type I | Worthington Biochemicals | LS004196 |
| <sup>18</sup>F-Fluorodeoxyglucose | Edinburgh Clinical Research Imaging Centre (Edinburgh, UK) | N/A |
| Insulin | Eli Lilly (Indianapolis, USA) | Humulin S |
| Ribozol™ | Amresco (USA) | N580 |
| Osmium Tetroxide | Agar Scientific (UK) | AGR1022 |
| OLIGONUCLEOTIDES |  |  |
| See Table 2 | This paper | N/A |
| EXPERIMENTAL MODELS: Organisms/Strains |  |  |
| Mouse: C57BL/6J | Charles River | 027 |
| Rabbit: New Zealand White | Envigo | HsdIf: NZW |
| SOFTWARE |  |  |
| Prism | GraphPad | v8.1.0 |
| Heatmapper | (Babicki et al., 2016) | http://www.heatmapper.ca/ |
| GSEA | (Subramanian et al., 2005). | v3.0 [build: 0160] |
| NRecon | Bruker microCT (Kontich, Belgium) | v1.6.9.4 |
| CT Analyzer | Bruker microCT (Kontich, Belgium) | v1.13.5.1 |
| PMOD | PMOD Technologies LLC (Zurich, Switzerland) | v3.806 |
| Analyze | AnalyzeDirect (Overland Park, KS, USA) | v12.0 |
| MatLab | Mathworks | R2018b |
| OTHER |  |  |
| TSE Phenomaster | TSE | PhenoMaster 1.0 with software version 6.1.9 |
| OneTouch Verio Glucometer | OneTouch | User's manual here |
| Gamma counter | PerkinElmer (USA) | Wizzard<sup>2</sup> |
| Preclinical PET/CT scanner | Mediso (Budapest, Hungary) | nanoScan PET/CT 122S |
| Clinical PET/CT scanner | Siemens Medical Systems (Erlangen, Germany) | Biograph mCT |
| MRI scanner | Siemens Medical Systems (Erlangen, Germany) | 3T Verio |
| Clinical CT scanner | Toshiba Medical Systems (Japan) | Aquilion ONE |

### Human subjects

For human subjects in cohort 1 (Fig. 1E, Supplemental Fig. 2A-B), ethical approval and subject characteristics are as described previously (Mattiucci et al., 2018). For human subjects in cohort 2 (Fig. 1F-G, Supplemental Fig. 2C-D) and those undergoing MRI or PET/CT (Fig. 4, Supplemental Fig. 5-6), all studies were reviewed and approved by the South East Scotland Research Ethics Committee, with informed consent obtained from each subject. Characteristics for cold-exposed, placebo-treated and prednisolone-treated subjects are as described previously (Ramage et al., 2016b). Characteristics for all other subjects are provided below in Table 1.

**Table 1.** Characteristics of subjects in human cohort 2 (*used for BMAd isolation and molecular analysis*), those who underwent paired CT and MRI analysis (*used to identify Hounsfield Units for BMAT*), and those without or with detectable BAT at room temperature (*Fig. 4, Supplemental Fig. 5-6*). Age and BMI are mean ± SD. *ND* = not determined.

| Study | Number of Subjects<br>(Male/Female) | Age | BMI | Diabetic | Osteoporotic |
| --- | --- | --- | --- | --- | --- |
| BMA*d* isolation | 10 (4/6) | 67.1 ± 5.9 | 31.6 ± 6.9 | 0% | 0% |
| MRI & CT | 33 (24/9) | 65.7 ± 8.1 | 29.1 ± 4.8 | ND | ND |
| Room temp<br>(no BAT) | 10 (2/8) | 51.5 ± 19.6 | 20.7 ± 2.4 | 0% | ND |
| Room temp<br>(active BAT) | 10 (2/8) | 51.1 ± 16.0 | 21.0 ± 2.2 | 0% | ND |

### Animals

Studies in New Zealand White rabbits were approved by the University of Michigan Committee on the Use and Care of Animals, with daily care overseen by the Unit for Laboratory Animal Medicine. Rabbit housing, monitoring and tissue isolation were done as described previously (Cawthorn et al., 2016). For cohort 1 (Fig. 1), male rabbits (3.14 ± 0.19 kg, mean ± SD) were fed a high-fiber diet (cat. No 5326, LabDiet), receiving 100 g/day (31.91 ±0.19 g/kg body mass/day; mean ±SD) until 22 weeks of age. For cohort 2 (Supplemental Fig. 1), male rabbits were fed the same high-fiber diet *ad libitum* (68.26 ± 4.82 g/kg body mass/day; mean ±SD) until 13 weeks of age. Rabbits in each cohort were then euthanized and tissues isolated for subsequent analysis.

Studies in C57BL/6JCrl mice were approved by the University of Edinburgh Animal Welfare and Ethical Review Board and were done under project licenses granted by the UK Home Office. Male C57BL/6J mice were bred in-house and housed on a 12 h light/dark cycle with free access to water and food, as indicated.

### Human cell and tissue isolation

For cohort 1, adipocytes were isolated from femoral head bone marrow (BMAds) or subcutaneous WAT (WAT Ads) as described previously (Mattiucci et al., 2018). For cohort 2, adipocytes from bone marrow, trabecular bone and WAT were isolated from patients undergoing hip-replacement surgery: BMAds were obtained from the proximal femoral diaphysis; trabecular bone adipocytes were from the proximal femoral metaphysis; and WAT Ads were from gluteofemoral subcutaneous WAT. Immediately after surgical isolation, tissues were washed and stored in ice-cold Dulbecco’s phosphate-buffered saline (DPBS, 14190250, Gibco) for transport to a sterile tissue culture hood. Therein, DPBS was decanted through a sterile 300 μm nylon filter to remove blood, lipid and small debris. The remaining washed tissue was then transferred to a sterile, pre-weighed petri dish (100 mm) and tissue mass recorded. A solution of collagenase type I (Worthington Biochemicals) was made at 1 mg/mL in Krebs-Ringer HEPES (KRH) buffer (120 mM NaCl, 2 mM KCl, 1 mM KH2PO4, 0.6 mM MgSO_4_, 1 mM CaCl_2_*2H_2_O, 82 mM HEPES, 5.5 mM D-Glucose, 1% BSA) pre-warmed to 37°C; sufficient volume was made to allow for 2 mL per mg tissue and the solution was passed through a 0.22 μm filter before use.

After weighing, each tissue was minced in the petri dish using a sterile scalpel and scissors, then transferred to a Falcon tube containing the collagenase solution. Tissues in collagenase were then incubated for 45 min in a shaking water bath (120 rpm) at 37°C. Next, collagenase-digested tissue was passed through a 300 μm nylon filter and the cells within the filtrate were washed with fresh KRH buffer. Samples were then centrifuged at 500 rcf for 5 min at 4°C. The floating adipocyte layer was transferred by pipette to a new tube to be used for RNA isolation; an aliquot was also analyzed histologically to confirm the presence of adipocytes. After aspirating and discarding the supernatant, the stromal vascular fraction (SVF) of cells within the pellet was resuspended in 2x volume of red blood cell lysis buffer (Cat. No. R7757, Sigma) and incubated at room temperature for 5 min to lyse erythrocytes. KRH buffer was added to bring the volume to 15 mL and samples were centrifuged at 700 rcf for 10 min at 4°C. The SVF pellet was then used for RNA isolation.

### RNA isolation and reverse transcription

For human cohort 1, RNA was extracted as described previously (Mattiucci et al., 2018). For human cohort 2 and mouse studies, RNA was isolated from cells or tissues using Ribozol™ solution (cat. No. N580, Amresco, USA,) according to the manufacturer’s protocol. For rabbit studies, RNA was isolated from tissues as described previously (Cawthorn et al., 2016). Tissues included iWAT, gWAT, dBMAT and ruBMAT of both cohorts, and pBMAT of rabbit cohort 2. RNA was quantified using a NanoDrop spectrophotometer (Thermo Scientific, USA), and cDNA was synthesized using the Taqman^®^ High Capacity cDNA Reverse Transcriptase Kit (Applied Biosystems, USA, cat. no. N8080234), in accordance with the manufacturer’s guidelines.

### Microarray analyses

For human cohort 1, RNA extraction, generation of single-strand biotinylated cDNA, and hybridization to Human GeneChip^®^ HTA 2.0 Arrays (Affymetrix) were done as described previously (Mattiucci et al., 2018). Transcripts were considered to be significantly differentially expressed between BMAds and WAT Ads when they had an adjusted *p*-value of 0.05 or less; *p*-values were adjusted for multiple comparisons using the Benjamini and Hochberg approach to control for false discovery rate (Benjamini and Hochberg, 1995).

For rabbit studies, purified RNA was digested on-column with DNase I and cleaned using the Qiagen RNeasy kit (Qiagen, Valencia, CA, USA) as recommended by the manufacturer. Total RNA was then submitted to the microarray core at the University of Michigan. The samples were screened for quality and processed in the microarray facility using custom rabbit Affymetrix arrays and the IVT Express kit (Affymetrix, Santa Clara, CA, USA). As a QC measure, the distribution of probe intensities and the 5’ to 3’ degradation profiles were checked to be consistent across samples. The core’s statistician used RMA, from the Affy package of Bioconductor, to fit log_2_ expression values to the data (Irizarry et al., 2003). Weighted, paired, linear models were then fit and contrast computed using the limma package (Smyth, 2004). Weighting was done using a gene-by-gene algorithm designed to down-weight chips that were deemed less reproducible (Ritchie et al., 2006). Probe-sets with a variance over all samples less than 0.05 were filtered out. Of the remainder, probe-sets with a log_2_-fold change of 2 or greater and an adjusted p-value of 0.05 or less were retained. P-values were adjusted for multiple comparisons using the Benjamini and Hochberg false discovery rate approach (Benjamini and Hochberg, 1995). Affy, affyPLM, and limma packages of Bioconductor, implemented in the R-statistical environment were used to analyze the data, including PCA analysis (Irizarry et al., 2003).

Pathways enriched in BMAT (combined dBMAT and ruBMAT) or WAT (combined iWAT and gWAT) of rabbits, or in isolated adipocytes from femoral BM and gluteofemoral scWAT (humans), were identified using Gene Set Enrichment Analysis (GSEA) software and the Molecular Signature Database (MSigDB) (Subramanian et al., 2005). For rabbits, pBMAT was not included in these analyses because it contained a high proportion of red marrow and therefore represented a less pure BMAT sample (Cawthorn et al., 2016). To ensure maximum compatibility with this software, rabbit gene identifiers were first converted to their corresponding human homologues using the BetterBunny algorithm (Craig et al., 2012). Volcano plots and heat maps (*Pearson Distance*) to visualize significantly differentially expressed transcripts (adjusted *p*-value <0.05, fold-change >2) were generated using Prism 8 (GraphPad) and Heatmapper software (Babicki et al., 2016), respectively.

### qPCR

For human cohort 2 and tissues from mice, reverse transcription, primer design/validation and qPCR were done as described previously (Sulston et al., 2016). Expression of each target gene was normalized to expression of 18S rRNA (human gene, *RNA18SN5;* mouse gene, *Rn18s), IPO8* or *Ppia*, based on consistency of housekeeper expression across all samples. For each transcript, expression is presented relative to the group with the highest expression. A Taqman assay was used to analyze *Ucp1* expression in mouse tissues (Thermo Fisher, Mm01244861_m1). All other primer sequences are described in Table 2.

**Table 2.**
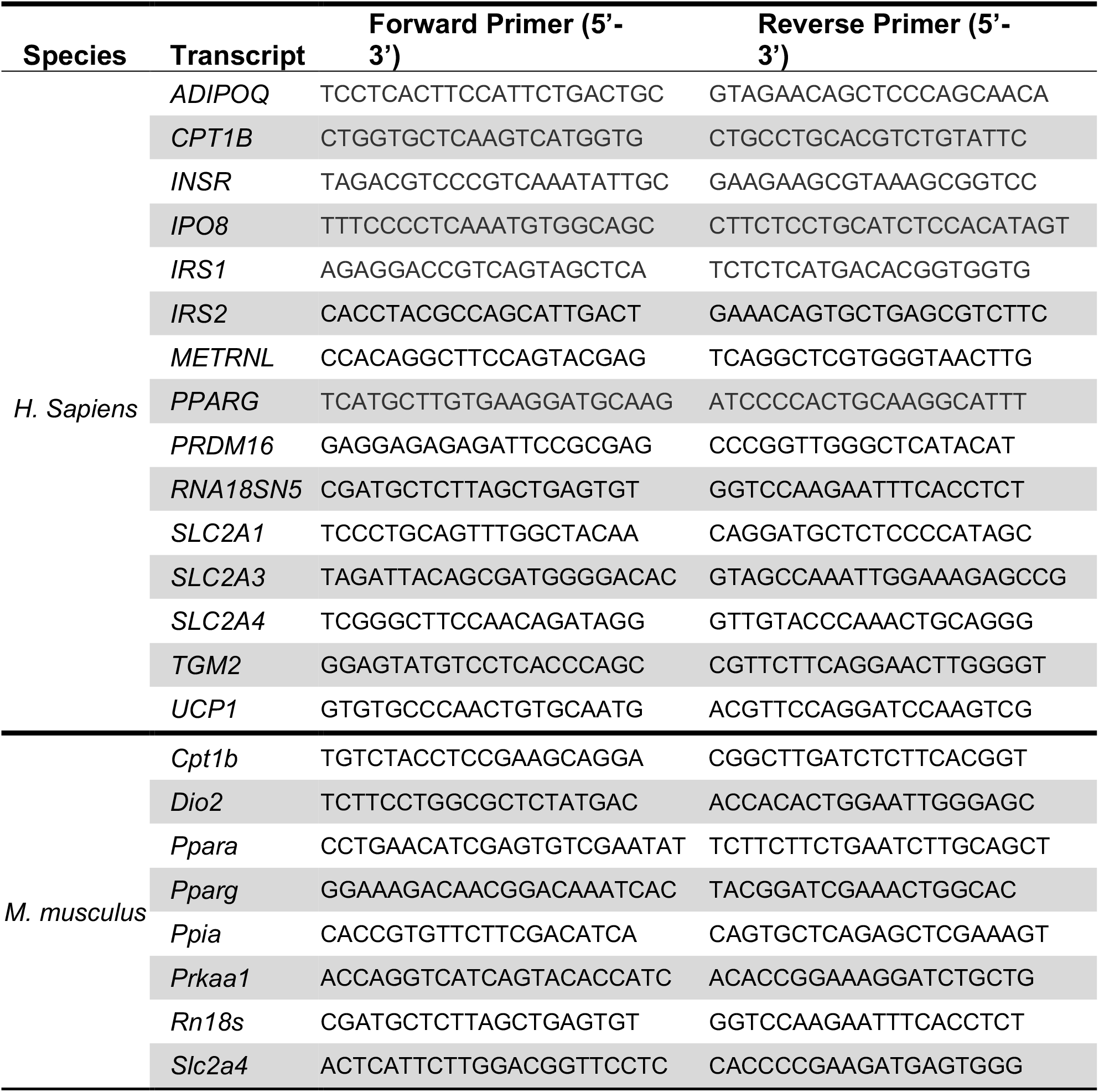
Sequences of primers used for qPCR.

### Histology

Fixed murine and human soft tissue and decalcified bones (14% EDTA for 14 days) were paraffin embedded by the histology core at The University of Edinburgh’s Shared University Research Facilities (SuRF). Paraffin-embedded tissue sections were then sectioned at 100 μm intervals using a Leica RM2125 RTS microtome and collected onto 76 x 26 mm StarFrost slides (VWR, UK). The slides were baked at 37°C overnight before Hemotoxylin and Eosin (H&E) staining.

### Mouse insulin-treatment studies

C57BL/6J male mice aged ~16 weeks were fasted for 4 h at room temperature (RT). Insulin (Humulin S, Eli Lilly; 0.75 mIU/g body mass) or sterile saline (0.9%) was then adminstred to to mice via intraperitoneal injection immediately prior to ^18^F-FDG injection. Mice were then returned to their cages. At 0 min (just before ^18^F-FDG injection), 15- and 60-min post-^18^F-FDG blood glucose was measured by tail venesection and blood sampled directly into EDTA-microtubes (Sarstedt, Leicester, UK). Mice were then anesthetised and ^18^F-FDG distribution assessed by PET/CT. After PET/CT, mice were sacrificed by overdose of anesthetic. BAT, iWAT, gWAT, pWAT, mWAT, gonads, brain, kidneys, liver, spleen, pancreata, heart, soleus, gastrocnemius, femur, tibiae, humeri, and tail vertebrae were then dissected and ^18^F-FDG uptake into each tissue was determined using a gamma counter (PerkinElmer). Counts per minute were converted to MBq activity using a standard conversion factor calibrated for the gamma counter. MBq were then corrected for radioactive decay based on the time of ^18^F-FDG administration and the time of gamma counting. Finally, the corrected MBq values were normalized to the mass of each tissue. The final gamma counts are therefore presented as *%* injected dose per g tissue (%ID/g). Frozen and fixed tissues were analyzed separately and the average MBq for each tissue was then calculated. Half of the dissected material was then snap frozen on dry ice and stored at −80°C for molecular analyses. The remaining half of the dissected material was placed into 10% formalin and stored at 4°C for histological analysis. PET/CT analysis was then done as described below.

### Mouse cold-exposure studies

The protocol is adapted from (Wang et al., 2012), with a summary depicted in Supplemental Figure 3A. For the acute and chronic cold exposure studies (Fig. 3, Supplemental Fig. 4B-H), male C57BL/6J mice aged ~18 weeks were housed individually in TSE PhenoMaster cages for indirect calorimetry, monitoring of physical activity, and measurement of *ad libitum* food and water consumption. Mice in each group were first housed in these cages for 3 days at room temperature (RT) for acclimation and baseline measurements. Group 1 (*Control*) were then housed for 72 h at RT in standard cages; group 2 (*Acute cold*) for 68 h at RT in standard cages, followed by 4 h at 4°C in TSE cages; and group 3 (*Chronic cold*) for 72 h at 4°C in TSE cages. Following TSE housing at 4°C, *Acute cold* and *Chronic cold* mice were returned to standard cages that had been pre-cooled on ice to 4 °C; *Control* mice continued to be housed in standard cages at RT. All groups were fasted, with access to water, for 4 h before administration of ^18^F-FDG (such that *Acute cold* mice were fasted throughout their 4 h cold exposure). Cages of cold-exposed mice were stored on ice in a ventilated cooler for transport to the PET/CT facility, while *Control* mice were transported at RT. After intraperitoneal injection of ^18^F-FDG, mice were returned to cages at RT (*Control*) or 4°C (*Acute* and *Chronic cold*). At 0 min (just before ^18^F-FDG injection), 15- and 60-min post-^18^F-FDG, blood glucose was measured by tail venesection. At 60-min post-^18^F-FDG, mice were placed under general anesthesia and underwent PET/CT imaging. Euthanasia, tissue isolation and gamma counting were done as described above for the insulin-treatment studies. PET/CT analysis was then done as described below.

To assess effects of cold exposure compared to mice housed at thermoneutrality, male C57BL/6J mice aged 12 weeks were individually housed for 48 h at 28°C, 22°C or 4°C. Each group was given AL access to chow diet throughout. Mice were then euthanised for tissue isolation.

### Mouse PET/CT analysis

PET/CT scan images were reconstructed and data was analyzed using PMOD version 3.806 (PMOD, Zurich, Switzerland). Standardised uptake values (SUV) were calculated for regions of interest, namely BAT, iWAT, gWAT; heart; bone tissue (without BM) from tibiae, femurs, and humeri; and the BM cavities within these bones. To distinguish bone tissue from BM, a calibration curve was generated using HU obtained from the acquisition of a CT tissue equivalent material (TEM) phantom (CIRS, model 091) and mouse CT scans. The TEM phantom consists of 2-4 mm hydroxyapatite rods representing mass densities of 1.08 to 1.57 g/mL. The TEM-reconstructed CT image data was exported for analysis into PMOD and, for the extraction of TEM HU values, a VOI template was generated and placed on each rod (0.008mL for 2mm and 0.05mL 4mm). The calibration curve was plotted based on the calculated linear equation of the TEM HU values, in which the mouse tissue values were inserted/scaled. This ensured that, within whole bones, regions of interest were specific for bone or BM.

### Human PET/CT studies

Subjects with active BAT at room temperature were identified retrospectively from clinical PET/CT scans; a control group, without detectable BAT, was then identified, ensuring that age, sex, weight, BMI and fasting blood glucose were matched to the active-BAT group. To assess effects of cold exposure, subjects were exposed to a mild cold (16 °C) for 2 h, as described previously (*Ramage et al., 2016b*). To assess effects of prednisolone treatment, subjects were recruited to a double-blind, randomized crossover study, as described previously (*Ramage et al., 2016b*). All subjects were placed supine in a hybrid PET/CT scanner (Biograph mCT, Siemens Medical Systems) and scanned as described previously (Ramage et al., 2016b).

### Determination of attenuation density for BMAT in humans

HU of subcutaneous fat, yellow marrow and red marrow were determined using Analyze 12.0 software (AnalyzeDirect, Overland Park, KS, USA) based on data from 33 patients who had undergone paired CT and MRI scans (Table 1). The MRI sequence was an axial HASTE (Half Acquisition Single Shot Turbo Spin Echo) with a TE (echo time) of 50 ms, TR (repetition time) of 1000 ms, and slice thickness 8mm. BM fat corresponds to higher signal intensity compared to surrounding bone and muscle tissues, in HASTE MR techniques. CT scanning was performed as described previously (Williams et al., 2017). Using Analyze 12.0 software, the MR and CT scans were co-registered and volumes of interest were manually drawn around the sternum, vertebrae and subcutaneous adipose tissue. HU were extracted on a per voxel basis, and data underwent post-processing using Matlab to measure the total number of voxels across all patient HU (Fig. 4B), prior to ROC analysis (MedCalc). ROC analysis was then conducted on per voxel HU to determine threshold values with the greatest sensitivity and specificity to detect bone, yellow marrow and red marrow. Thresholds of above 300 HU were defined as bone regions, −200 to 115 HU as yellow marrow and 115 to 300 as red marrow.

### Micro-computed tomography scanning (μCT)

Following euthanasia, murine tibiae were isolated, thoroughly cleaned and fixed in 10% formalin at 4°C for 48 hours. Bones were decalcified for 14 days in 14% EDTA and washed in Sorensen’s phosphate buffer. Bones were then stained for 48 hours in 1% osmium tetroxide (Agar Scientific, UK), washed in Sorensen’s phosphate buffer and embedded in 1% agarose, forming layers of five tibiae arranged in parallel in a 30-mL universal tube. Tubes of embedded tibiae were then mounted in a Skyscan 1172 desktop micro CT (Bruker microCT, Kontich, Belgium). Samples were scanned through 360° using a step of 0.40° between exposures. A voxel resolution of 12.05 μm was obtained in the scans using the following control settings: 54 kV source voltage, 185 μA source current with an exposure time of 885 ms. A 0.5 mm aluminum filter and two-frame averaging were used to optimize the scan. After scanning, the data were reconstructed using NRecon v1.6.9.4 software (Bruker, Kontich, Belgium). The reconstruction thresholding window was optimized to encapsulate the target image. Volumetric analysis was performed using CT Analyzer v1.13.5.1 (Bruker microCT, Kontich, Belgium).

### Statistical analysis

Microarray data were analyzed as described above. All other data were analyzed for normal distribution using the Shapiro-Wilk normality test. Normally distributed data were analyzed by ANOVA or t-tests, as appropriate. Where data were not normally distributed, non-parametric analyses were used. When appropriate, *P* values were adjusted for multiple comparisons. Data are presented as histograms or box and whisker plots. For the latter, boxes indicate the 25th and 75th percentiles; whiskers display the range; and horizontal lines in each box represent the median. All statistical analyses were performed using Prism software (GraphPad, USA). A *P*-value <0.05 was considered statistically significant.

## SUPPLEMENTAL FIGURE LEGENDS

**Figure S1, Related to Figure 1 –.**
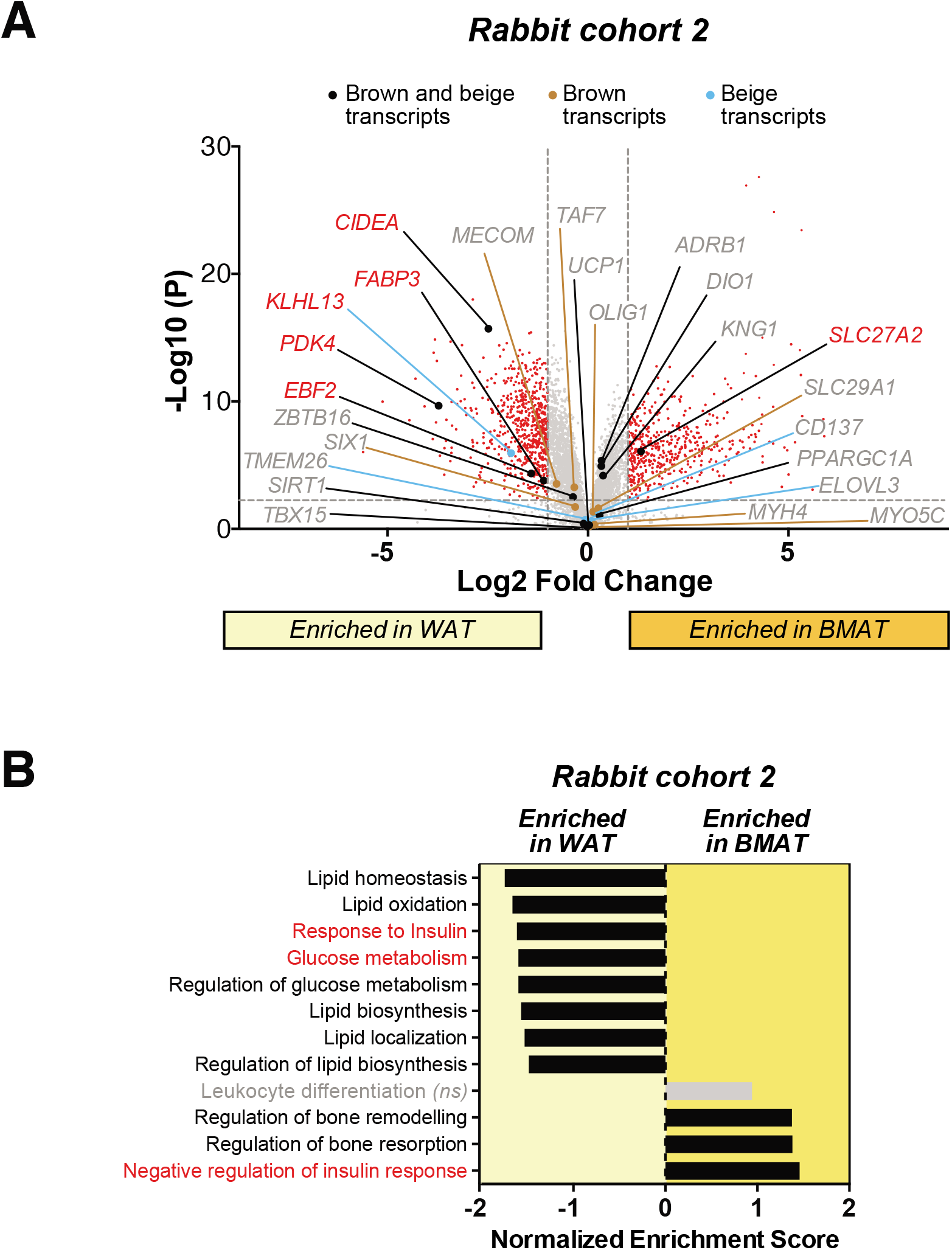
BMAT is transcriptionally distinct to white, brown and beige adipose tissues. Transcriptional profiling of WAT (gonadal + inguinal) vs whole BMAT (dBMAT + ruBMAT) from rabbit cohort 2. **(A)** Volcano plot of differentially expressed transcripts (FDR < 0.05, fold-change > 2). Transcripts characteristic of brown and/or beige adipocytes are labelled. Those with significant differential expression between WAT and BMAT are shown in red; those not differentially expressed are in grey. **(B)** Gene set enrichment analysis (GSEA) highlights lipid metabolism, glucose metabolism and insulin responsiveness as key pathways differentially regulated between WAT and BMAT.

**Figure S2, Related to Figure 1 –.**
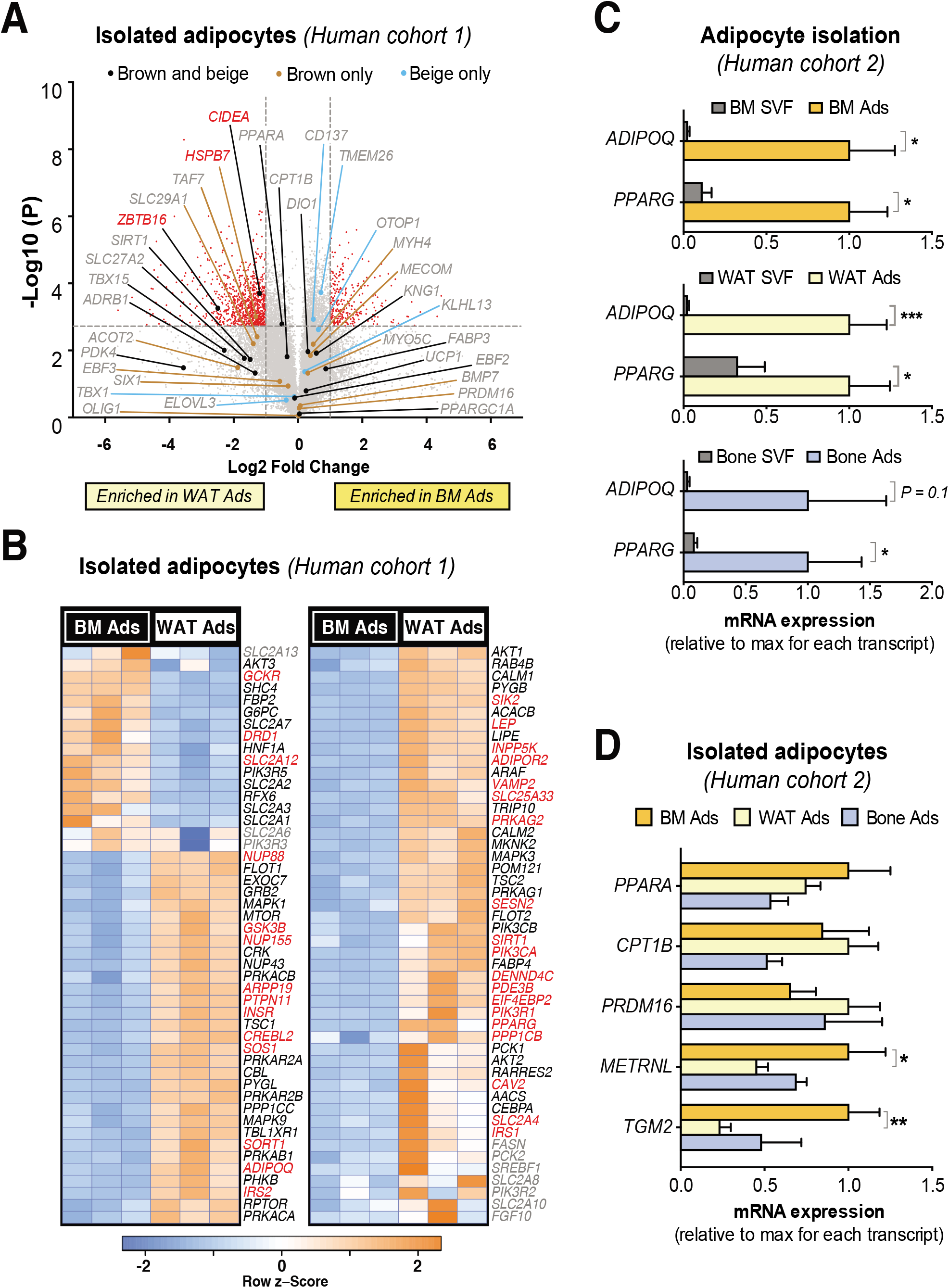
BM adipocytes in humans are transcriptionally distinct to those from WAT. Transcriptional profiling (A,B) and qPCR analysis (C,D) of adipocytes isolated from the subcutaneous WAT or femoral diaphyseal BM of humans undergoing hip-replacement surgery. **(A,B)** Volcano plots (A) and heat maps (B) are presented as for Figures 1 and S1. **(C,D)** qPCR to validate purity of adipocytes isolated from each tissue (C) and showing that BM adipocytes generally do not have increased expression of brown or beige adipocyte markers (D). Transcript expression was normalised to expression of *IPO8* (C) or *RNA18SN5* (D). Data in (C) are mean ± SEM of the following numbers per group: BM Ads, n = 8 (*ADIPOQ*) or 9 (*PPARG);* BM SVF, n = 3 (*ADIPOQ*) or 5 (*PPARG);* WAT Ads, n = 10 (*ADIPOQ* and *PPARG);* WAT SVF, n = 5 (*ADIPOQ* and *PPARG);* Bone Ads and SVF, n = 7 (*PPARG*) or 3 (*ADIPOQ*). Data in (D) are mean ± SEM of the following numbers per group: BM Ads, n = 3-10; WAT Ads, n = 6-10; Bone Ads, n = 3-7. For each transcript, significant differences are indicated as for Figure 1.

**Figure S3, Related to Figure 3 –.**
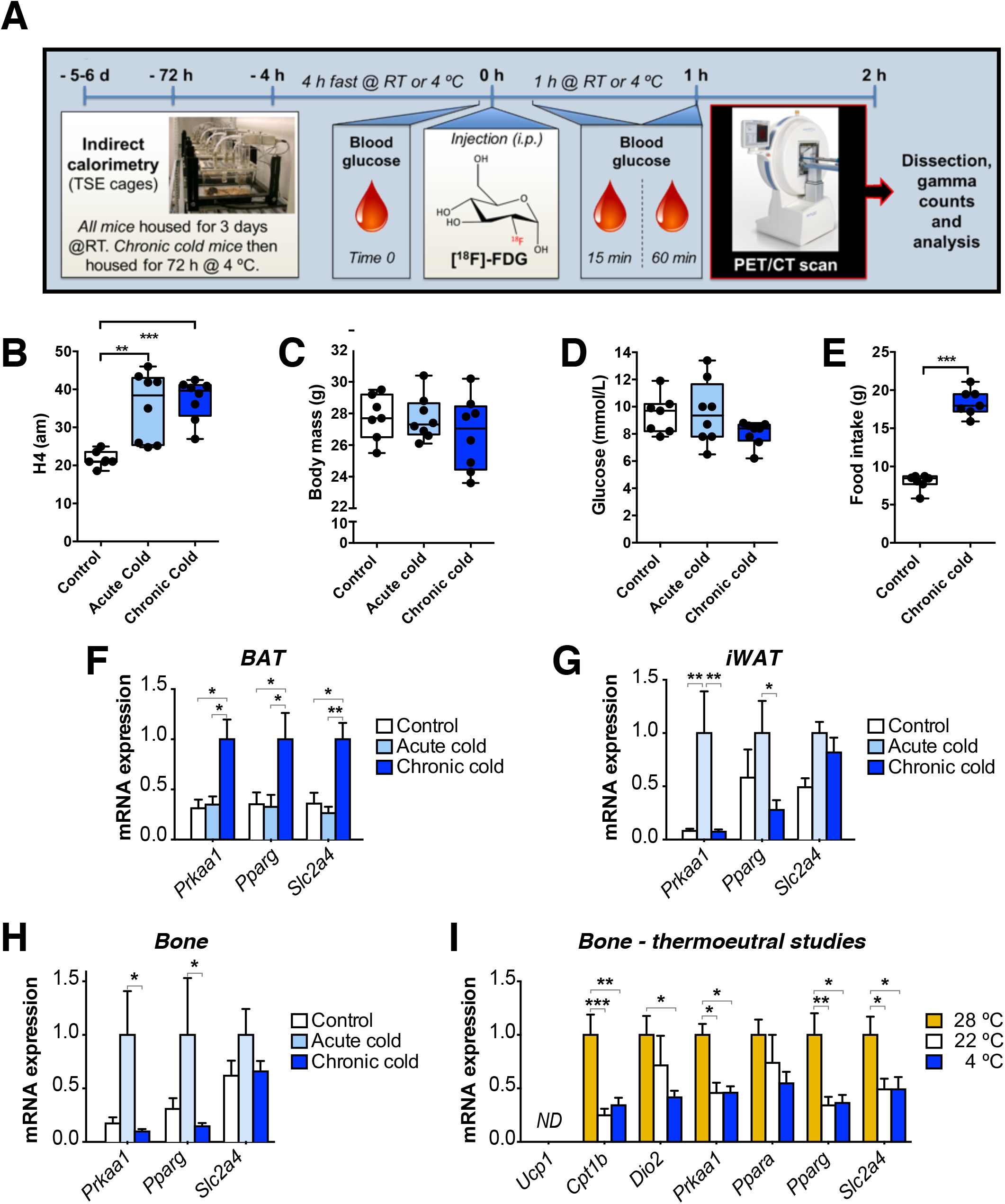
Effects of cold exposure on energy homeostasis and gene expression in BAT, iWAT and bone. **(A)** Protocol for cold exposure and calorimetry studies, as described in the STAR Methods. **(B-H)** Effects of cold exposure on energy expenditure (B), body mass (C), baseline blood glucose (D), 72 h food intake (E) and transcript expression in BAT, iWAT or whole femurs (F-H). In (E), *Acute cold* mice are not shown because they were fasted throughout cold exposure. **(I)** A separate cohort of mice was housed at thermoneutrality, 22 °C or 4 °C for 48 h. Expression of BAT or beige cold exposure markers was then determined by qPCR of whole femurs. *ND =* not detectable. In (F-I), transcript expression was normalised to expression of *Rn18s* (F,H,I) or *Ppia* (G); the latter was used for iWAT because in this tissue *Rn18s*, but not *Ppia*, showed significant regulation between the three groups. Data are shown as box-and-whisker plots (B-E) or as mean ± SEM (F-H) of 7-8 mice per group. In (I), data are mean ± SEM of 8-10 mice per group. Significant differences between groups are indicated by * (*P* <0.05), ** (*P* <0.01) or *** (*P* <0.001).

**Figure S4, Related to Figure 4 –.**
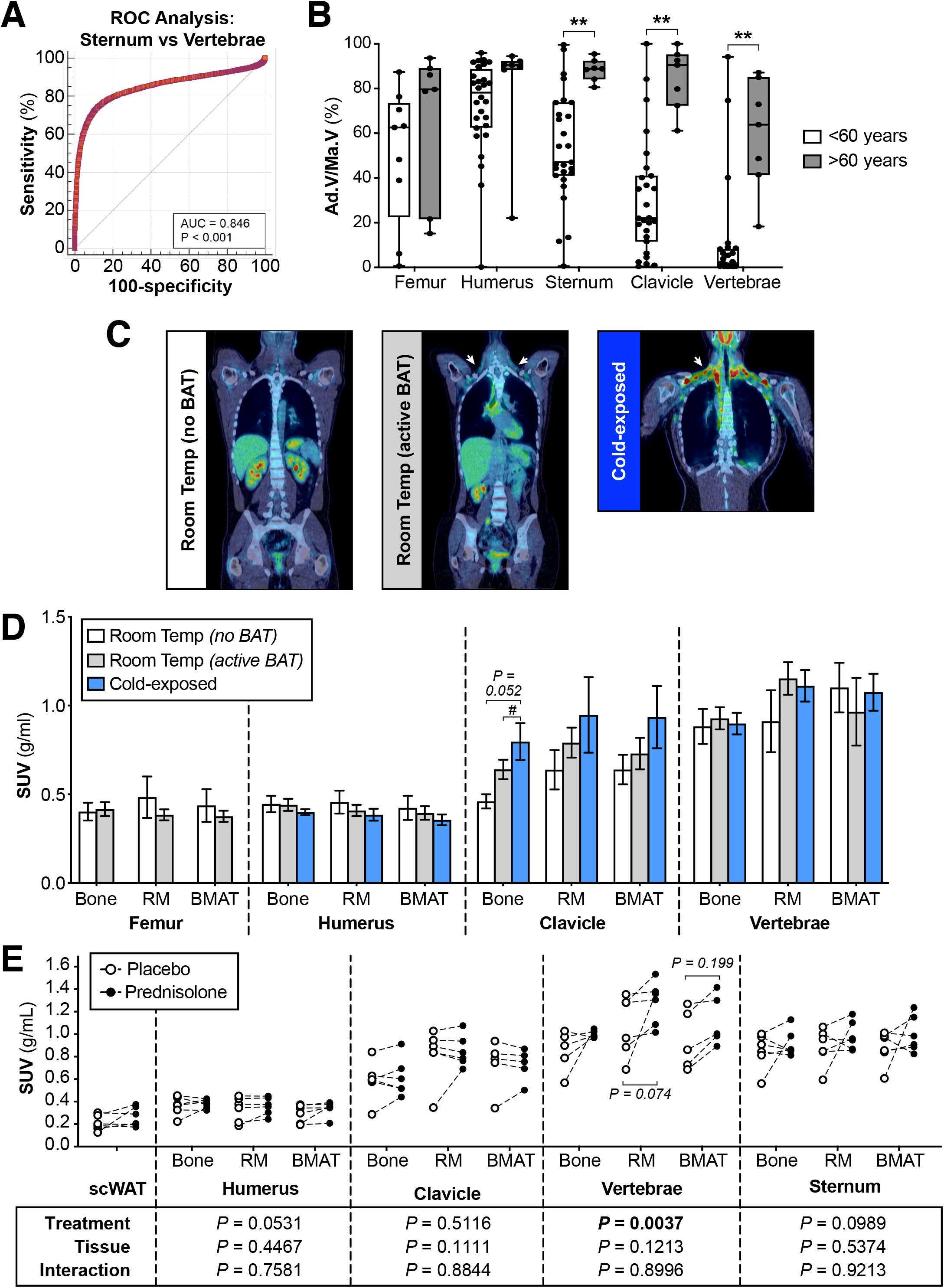
PET/CT for identification and functional analysis of BMAT in humans. **(A)** ROC analysis to identify HU thresholds to distinguish BMAT-rich (sternum) from BMAT-deficient (vertebrae) regions of BM. **(B)** Quantification of BMAT in CT scans of male and female subjects aged <60 or >60 years. A HU threshold of <115 was used to identify BMAT voxels in BM of the indicated bones, and total BM volume was also determined. The proportion of the BM cavity corresponding to BMAT (Ad.V/Ma.V) was then calculated. Data are shown as box-and-whisker plots of the following numbers of subjects for each group: <60 years, n = 28 (humerus), 9 (femur), or 27 (clavicle, sternum and vertebrae); >60 years, n = 7 for each bone. Significant differences between <60 and >60 groups are indicated by ** (*P* <0.01). **(C)** Representative coronal PET/CT images of *No BAT, Active BAT* and *Cold* subjects. ^18^F-FDG uptake in BAT is evident in the *Active BAT* and *Cold* subjects (arrows). Femurs were not included in scans of the *Cold* group. **(D)** ^18^F-FDG uptake in bone tissue, RM and BMAT of the indicated bones. Data are shown as mean ± SEM of 8 (*No BAT*) or 7 (7 *Active BAT, Cold*) subjects per group. Significant differences between these groups *No BAT, Active BAT* and *Cold* groups are indicated by # (*P* <0.05). **(E)** Subjects were treated with prednisolone or placebo control prior to analysis of ^18^F-FDG uptake by PET/CT. Data are shown as paired individual values for each subject. For each skeletal site, the influence of treatment or tissue (bone, RM, BMAT), and interactions between these, were determined by 2-way ANOVA; *P* values are shown beneath the graph.

